# Dynamically expressed ELAV is required for learning and memory in bees

**DOI:** 10.1101/2021.06.24.449637

**Authors:** Pinar Ustaoglu, Jatinder Kaur Gill, Nicolas Doubovetzky, Irmgard U. Haussmann, Jean-Marc Devaud, Matthias Soller

## Abstract

Changes in gene expression are a hallmark of learning and memory consolidation. Little is known about how alternative mRNA processing, particularly abundant in neuron-specific genes, contributes to these processes. Prototype RNA binding proteins of the neuronally expressed ELAV/Hu family are candidates for roles in learning and memory, but their capacity to cross-regulate and take over each other’s functions complicate substantiation of such links. Therefore, we focused on honey bees, which have only a single *elav* family gene. We find that honey bee *elav* contains a microexon, which is evolutionary conserved between invertebrates and humans. RNAi knockdown of *elav* demonstrates that ELAV is required for learning and memory in bees. Indicative of a role as immediate early gene, ELAV is dynamically expressed with altered alternative splicing and subcellular localization in mushroom bodies, but not in other brain parts. Expression and alternative splicing of *elav* change during memory consolidation illustrating an alternative mRNA processing program as part of a local gene expression response underlying memory formation. Although the honey bee genome encodes only a single *elav* gene, functional diversification is achieved by alternative splicing.

## Introduction

Changes in gene expression play pivotal roles in memory consolidation, the process through which memories are stabilized and stored into long-term memory ^1-3^. Among genes induced upon learning are immediate early genes (IEGs) such as transcription factors *egr-1, fos* and *jun*, and changes in transcription seems to be a highly conserved requirement for memory formation, including in insects ^4-10^. A common feature of neuronal genes, particularly ion channel and cell adhesion genes, is their often complex pattern of alternative splicing, which alters protein coding and regulatory potential in flanking untranslated regions of the mRNA ^11-13^. Since very little is known about alternative splicing programs that operate in learning and memory we focused on ELAV (Embryonic Lethal Abnormal Visual system)/Hu family RNA binding proteins because they are prominently expressed in neurons of all metazoans, regulate alternative splicing and expression of synaptic genes as well as formation of new connections ^14-18^.

Like many RNA binding proteins (RBPs) ELAV/Hu proteins comprise a family of highly related proteins in most animals including humans. Humans have four *ELAV/Hu* genes (*HuB, HuC, HuD* and *HuR*), while *Drosophila* have three (*elav, fne* and *Rbp9*), which derive from a common ancestor, but have duplicated independently during vertebrate and arthropod evolution ^19^. In mice, all Hu proteins are expressed in largely overlapping patterns in mature neurons ^20^, while in *Drosophila* pan-neural expression of ELAV and FNE starts with the birth of neurons, and RBP9 is first detected in late larval neurons ^21-24^. Although ELAV family RBPs in *Drosophila* have distinct neuronal phenotypes based on the analysis of null mutants and genetic interactions among them, they can cross regulate each other’s targets depending on cellular localization and concentrations complicating the analysis of their functions ^24^.

ELAV/Hu proteins are proto-type RBPs, which harbor three highly conserved RNA Recognition Motifs (RRMs). The first two RRMs are arranged in tandem and the third RRM is separated by a less-conserved hinge region. ELAV/Hu family RBPs bind to short, uridine-rich motives, which are ubiquitously found in introns and untranslated regions, but ELAV/Hu proteins are gene-specific and have a complement of dedicated target genes ^18,25-27^. Due to the prominent nuclear localization, ELAV in *Drosophila* has mostly been associated with gene-specific regulation of alternative splicing and polyadenylation, but it can also regulate mRNA stability ^28-34^. Although the three RRMs comprise the evolutionary most conserved parts of ELAV/Hu proteins, individual members are to a large degree functionally interchangeable when adjusting expression levels and sub-cellular localization ^24,35,36^. Hence, regulation of the activity of ELAV/Hu proteins likely occurs at the level of post-translational modifications and suggest that less conserved and unstructured linker sequences between or within RRMs serve fundamental functional roles, possibly by regulating interactions with other proteins ^37^.

To avoid complications of assigning specific gene functions to individual members of the ELAV/Hu family, we focused on honey bees whose genome encodes only one copy of the *elav* gene ^19^, an orthologue of *Drosophila fne* ^22^. Conveniently, honey bees are a well-established model for the study of learning and memory. Here we show that the single *elav* gene in honey bees is required for learning as well as formation of stable memories by RNAi knockdown. Although bees have only a single *elav* gene, its coding capacity proliferated by increasing alternative splicing to generate 38 different isoforms. The splicing pattern changes during development and between different adult social castes, but also shows variability among brains of individua adult workers. Likewise, ELAV expression changes in mushroom bodies (brain centers involved in learning and memory), but not in the medulla of the optic system, to generate individual expression patterns reminiscent of experience dependent neuronal activity that forms the basis of gene expression changes associated with memory consolidation. Hence, ELAV expression resembles that of immediate early genes (IFGs) induced upon different experiences ^38^. Consistent with a role in learning and memory consolidation, *elav* expression and inclusion levels of alternative exons change during the early phases of memory consolidation that requires transcription ^10,39^.

## Results

### ELAV is required for learning and memory consolidation in bees after olfactory reward conditioning

To assess whether ELAV has a role in learning and memory in bees the single bee *elav* gene was knocked down by RNAi leading to a reduction of 80% after two days (n=3, Fig 1A). Two days after injection of *elav* or *GFP* control dsRNA, bees were individually trained and short-term memory was scored 2 hours after training (Fig 1B). Both groups showed significant learning over the successive trials (RM-ANOVA, *Trial* effect: F=61.93, p<0.001), but performance was affected by treatment (*Trial* x *Treatment* interaction: F=4.33, p<0.05). Indeed, as compared to controls, significantly fewer *elav* dsRNA-injected bees showed conditioned responses by the end of training (Fischer’s test on 3^rd^ trial: χ^2^ = 4.22, p<0.05, Fig 1C left). However, short-term memory retrieval remained unaffected (χ^2^ = 0.64, p>0.05, Fig 1C right).

**Figure 1.**
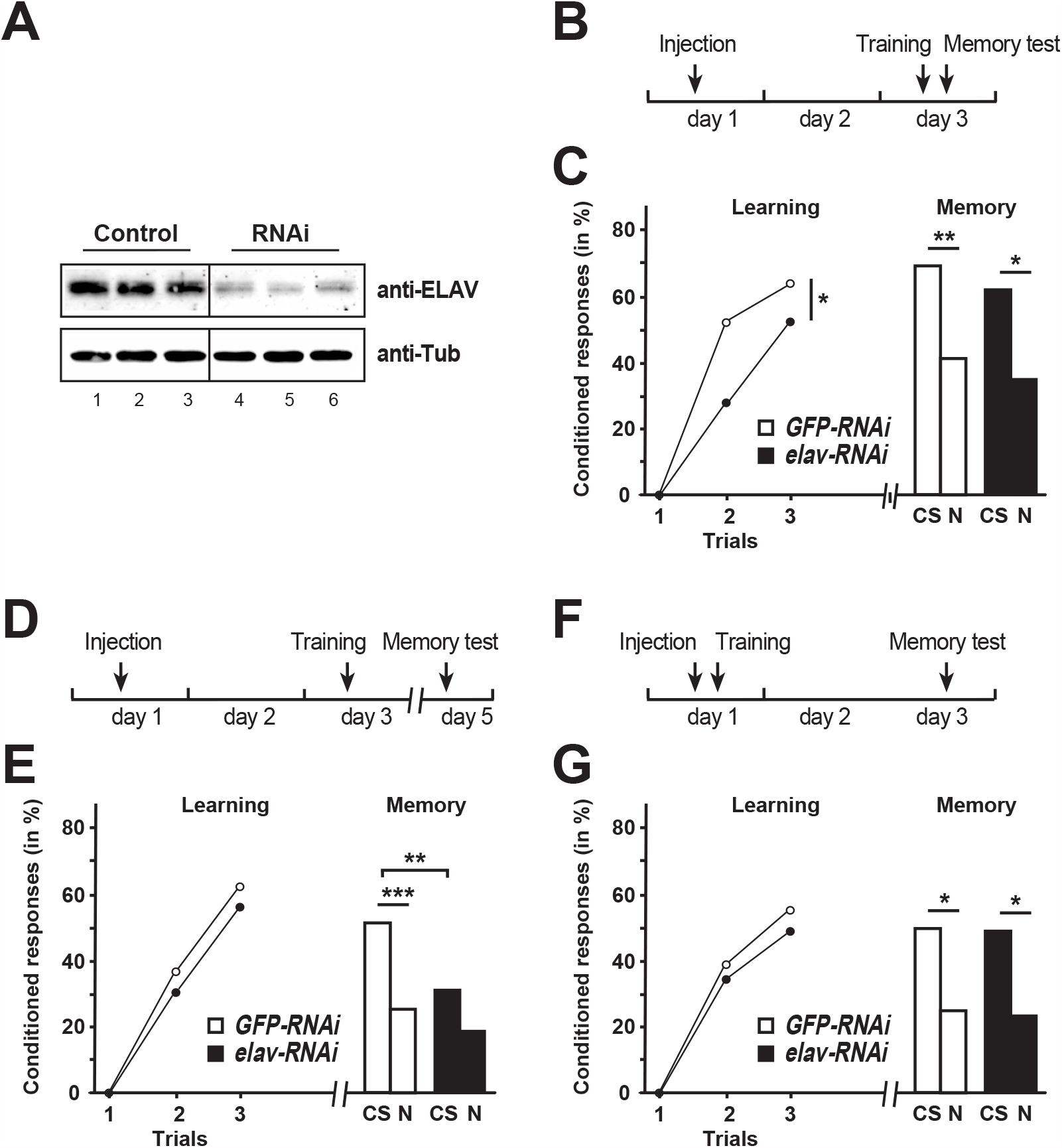
Fig 1: ELAV is required for learning and memory consolidation. (A) Western blot detecting ELAV in bee central brains of control *GFP* and *elav* dsRNA injected workers 50 h after injection. (B) Schematic of the treatment to test for ELAV’s role in learning. (C) Learning (*left*) and memory (*right*) performances of control *GFP* dsRNA (*white*, n=66) and *elav* dsRNA (*black*, n=74) injected worker bees. *CS*: conditioned odor, *N*: novel odorant. *: p<0.05; **: p<0.01. (D) Schematic of the treatment to test for ELAV’s role in memory consolidation. (E) Learning (*left*) and memory (*right*) performances of control *GFP* dsRNA (*white*, n=74) and *elav* dsRNA (*black*, n=77) injected worker bees. *CS*: conditioned stimulus, *N*: novel odorant **: p<0.01; ***: p<0.001. (F) Schematic of the treatment to test for ELAV’s role in memory retrieval. (G) Learning (*left*) and memory (*right*) performances of control *GFP* dsRNA (*white*, n=53) and *elav* dsRNA (*black*, n=50) injected worker bees. *CS*: conditioned stimulus, *N*: novel odorant *: p<0.05.

We then asked whether *elav* knockdown might impact on the consolidation of long-term memory independently on its effect on acquisition. Therefore, injections and training were performed as before to ensure that *elav* levels would still be reduced during the hours following training (Fig 1D), i.e. at a time when crucial transcriptional activity is required for long-term memory consolidation ^10,39^. We then tested for their memory two days after training (a typical delay to assess consolidated long-term memory). In these conditions, learning occurred normally (RM-ANOVA, *Trial* effect: F=108.6, p<0.001; *Trial* x *Treatment* interaction: F=0.50, p>0.05; Fig 1E left). Yet, the two groups showed different capacities to recall the memory of the CS-US association (Fischer’s test: χ^2^ = 10.08, p<0.01, Fig 1E right). In addition, only control bees responded significantly more to the CS than to the novel odorant (*GFP*: χ^2^ = 11.55, p<0.001; *elav*: χ^2^ = 3.77, p>0.05).

To reject the possibility that loss of ELAV impairs long-term memory retrieval *per se* due to a prolonged downregulation of *elav*, we performed an additional experiment in which injection was done shortly before training, when RNAi is not yet effective (Fig. 1F). As expected, this treatment did not affect learning (*Trial* effect: F=62.93, p<0.001; *Trial* x *Treatment* interaction: F=0.15, p>0.05; Fig. 1G left). More importantly, memory retrieval was intact and two days after training both groups responded similarly to the CS (Fischer’s test: χ^2^ = 0.02, p>0.05) and responded significantly less to the novel odorant (*GFP*: χ^2^ = 6.24, p<0.05; *elav*: χ^2^ = 5.66, p<0.05), thus indicating a preserved memory of the CS-US association.

These results thus argue that *elav* is required for the early formation of an associative memory over repeated acquisition trials, and for its subsequent consolidation.

### The single bee ELAV gene is dynamically alternatively spliced

The bee ELAV protein is highly homologous to those of the *Drosophila* ELAV family (ELAV, FNE and RBP9) in the three RRM domains, but diverges significantly in the unstructured hinge domain separating RRM2 from RRM3 (Fig 2A and Supplemental Fig S1A). Given the much more sophisticated tasks associated with the social life of bees, the presence of only single ELAV in their brains, compared to three pan-neuronally expressed genes in brains of adult flies, was surprising and prompted us to investigate whether *elav* in bees is alternatively spliced to compensate for the lack of the expected genic diversity. Indeed, cloning of full-length *elav* from RT-PCR revealed five alternatively spliced exons: exons 3a, exon 4a adding an additional 3’ss, exon 4b adding an additional 5’ss, exon 4c and exon 4d, (Fig 2A-C, Supplemental Fig S1). The combination of these exons in addition to skipping of exon 4 variables potentially generates 38 different isoforms (Fig 2A).

**Figure 2.**
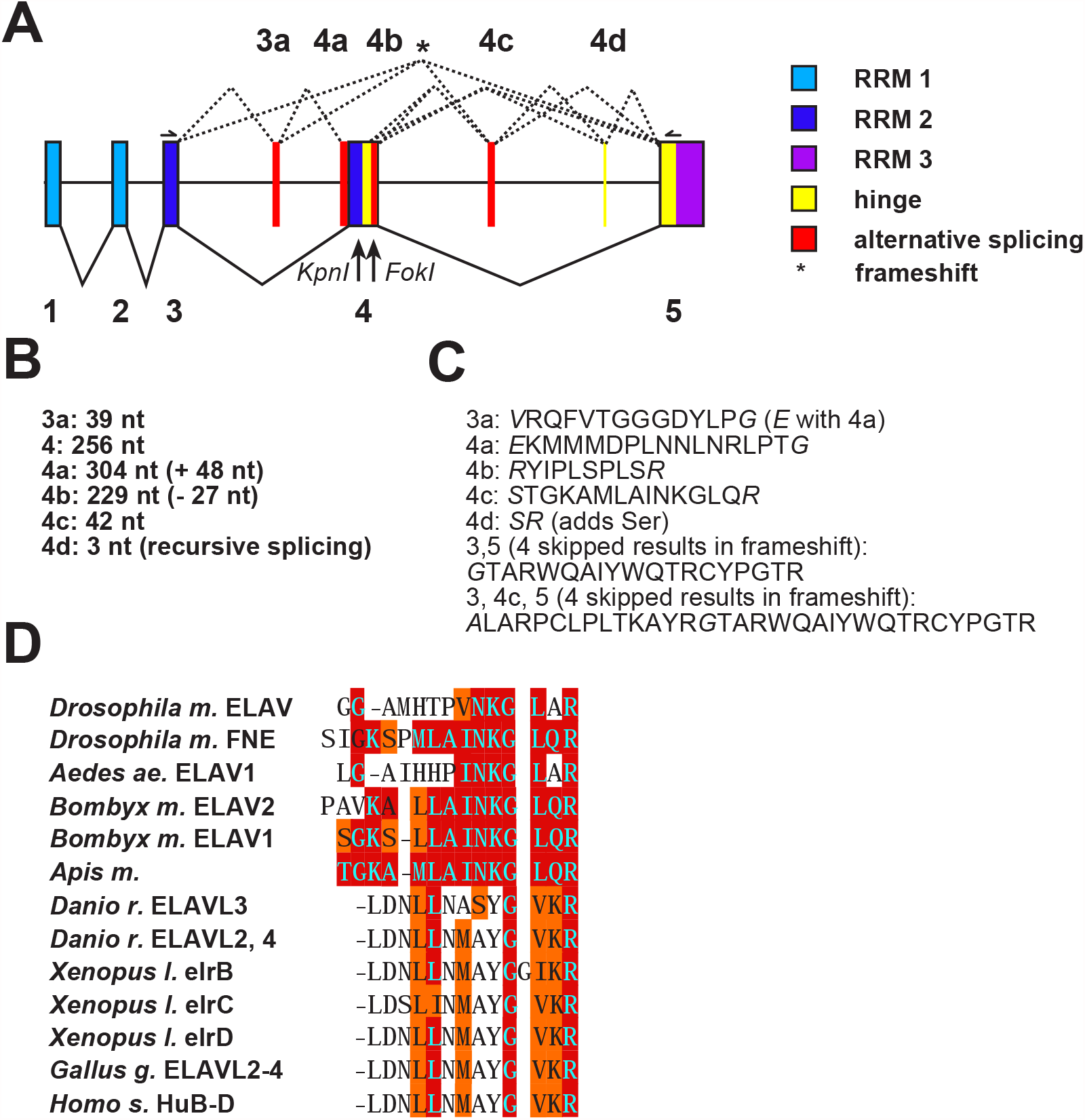
Single Bee *elav* gene is alternatively spliced. (A) Gene model of *Apis mellifera elav* gene depicting exons (boxes) and their splicing. In total, 38 different alternative splice products are possible. (B) Length of alternative microexons. (C) Sequence of alternative microexons. (D) Alignment of alternative microexon 4c from bees with a part present in *Drosophila* ELAV or alternatively spliced microexons in ELAV/Hu family proteins of other species. Note that *Apis* ELAV microexon 4c is the first alternatively spliced exon shown to be conserved between vertebrates and insects.

Intriguingly, two of these alternative exons are located in the loop region of RRM2 and the other three are located in the hinge region (Fig 2A-C, Supplemental Fig S2A). Exon 4d is only 3 nt long and codes for a serine which potentially can be phosphorylated to impose further control of ELAV function ^37^. Since the sequence of exon 4d is TAG and flanked by AG/GT consensus splice sites it is not a substrate for recursive splicing ^40-42^. Exon 4d is too small to accommodate spliceosomal complexes on both sides and must thus be spliced sequentially ^11^.

Rather unexpectedly, we also detected isoforms which skip exon 4 and its variants encoding the second half of RRM2. This results in truncated ELAV proteins by introducing a frameshift removing much of the beta-sheet of RRM2 involved in RNA recognition as well as alpha-helix 2 that makes up the backbone of the RRM structure. Since skipping of variable exons between exon 3 and 4 deemed unfunctional based on RNA binding assays ^43^, we employed molecular modeling to explore the capacity of frequently included alternative exons 3a and 4a to build alternative structures that might hold functionality. Indeed, inclusion of exon 3a with concomitant exclusion of exon 4, 4a and/or 4b adds an additional beta-sheet potentially increasing the capacity to bind RNA (Supplemental Fig 2B). Inclusion of exon 4c further adds an additional alpha-helix likely stabilizing this alternative RRM structure (Supplemental Fig 2C).

Intriguingly, exon 4c from bees has been found conserved in *Drosophila* FNE and aligns to part of ELAV ^35,36,44^. Human ELAV/Hu family proteins also harbor an alternatively spliced small exon between the second and third RRM at a position similar to that of the hinge region of bee *elav*, before a conserved motif involved in nuclear cytoplasmic shuttling ^45^. Alignment of exon 4c from bees with orthologues in other insects (the mosquito *Aedes aegyptii* and the moth *Bombyx mori*), zebrafish *Danio rerio*, African clawed frog *Xenopus laevis*, Chicken *Gallus gallus* and human ELAV/Hu proteins revealed evolutionary conservation of this microexon (Fig 2D), which is consistent with an evolutionary conserved microexon program between vertebrates and invertebrates ^46,47^.

Bees also have a 3 nt microexon (Fig 2A-C), that adds a serine, which potentially can be phosphorylated ^37^. In vertebrates, this serine is added through an alternative 3’splice site at the same position in Human *HuB* and *HuC*, chicken *ELAVL2-4*, Xenopus *elrB* and *elrC*, and zebrafish *ELAVL2* and *ELAVL3*.

Next, we analyzed alternative splicing in more detail than possible on agarose gels, where multiple alternative splice products, amplified from mRNA of larval brains, where detected only as a smear (Fig 3A). Consistently with the small differences of alternative splice products, alternative protein isoforms were not separable either by Western blot detection (Fig 3B). Therefore, we employed a higher resolution separation of ^32^P-labeled PCR products using denaturing polyacrylamide gels. This analysis revealed 23 distinguishable products with sizes between 78 and 463 nt (Fig 3 C and D). Most frequently found isoforms were 3-4-5, 3-3a-4-4c-5, 3-3a-4-5/3-4-4c-5 and 3-3a-4b-5/3-4b-4c-5 as well as the truncated isoforms 3-4c-5 and 3-5, thus indicating functional relevance for the newly identified alternative splice products.

**Figure 3.**
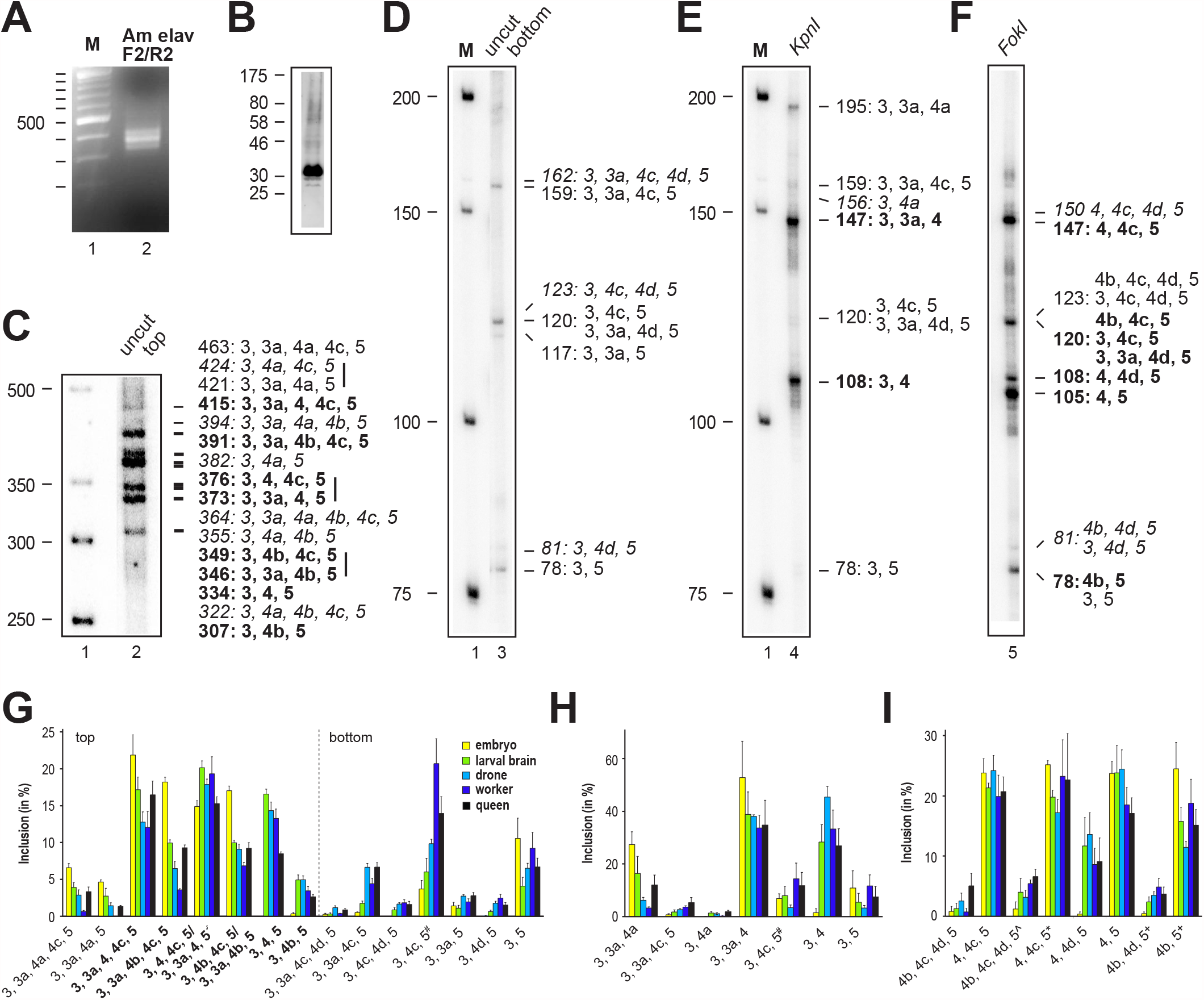
*elav* alternative splicing is dynamic during development. (A) Multiple products are detected from RT-PCR of RNA from larval brain in the alternatively spliced part of bee *elav* on an agarose gel. M: Marker (B) Western blot of bee ELAV in larval brains. (C and D) Top and bottom part of a representative 5 % denaturing polyacrylamide gel separating ^32^P-labeled alternative splice products from larval brains. Length of PCR products from splice variants are indicated at the right in bold for prominent products and in italics for very rare products. Vertical lines at the right of indicated splice variants indicate inseparable products. M: Marker. (E and F) Analysis of alternative splicing from larval brains proximal (*Kpn*I) and distal (*Fok*I) of exon 4 from ^32^P-labeled labeled forward (E) or return (F) primer after digestion with either *Kpn*I or *Fok*I on a representative 5 % denaturing polyacrylamide gels. Length of PCR products from splice variants are indicated at the right in bold for prominent products and in italics for very rare products. M: Marker. (G-I) Developmental and sex-specific alternative splicing of bee *elav* quantified from denaturing polyacrylamide gels shown as mean with the standard error from three replicates as percent from all splice products from top (G left) and bottom (G right) gel parts and after *Kpn*I or *Fok*I digestion as above from embryos (yellow), larval brains (green), drone brains (light blue), worker brains (dark blue) and queen brains (black). (G and H) #: The 120 nt product can be either 3, 4c, 5 or 3, 3a, 4d, 5. (I) ^: The 123 nt products are either 4b, 4c, 4d, 5 or 3, 4c, 4d, 5. *: The 120 nt products are either 4b, 4c, 5 or 3, 4c, 5 or 3, 3a, 4d, 5. ^+^: The 80 nt products are either 4b, 5 or 3, 5 +/- 4d.

Since some of the isoforms were not separable based on size, we wanted to determine how frequently each alternative exon is included. For this purpose, we digested 5’ ^32^P-labeled PCR products with KpnI or FokI restriction endonucleases to cleave off their unlabeled 3’ parts (Fig 2C, 3E and 3F). For both sides of exon 4, all possible combinations of alternative splice products were detected.

Next, we analyzed the ELAV alternative splicing pattern at different developmental stages and in different tissues (n=3, Fig 3G-I and Supplemental Fig 3). This analysis revealed dynamic inclusion of alternative exons. Most strikingly, splicing from exon 3 to 4 is absent in embryos and skipping of exon 4, 4a or 4b leads to considerably increased abundance of the truncated isoform 3-4c-5 in adults, particularly workers.

To obtain further insights into the dynamics of *elav* alternative exon use at a cellular level we performed whole mount RNA in situ hybridization with anti-sense probes against alternative exons 3a and 4c in brains of worker bees. Most strikingly, both exons 3a and 4c show very dynamic inclusion levels in the mushroom bodies, displaying unique patterns in each individual bee (Fig 4 A, D, J and M). In contrast, inclusion levels in the medulla (visual neuropile not involved in the learning process) are uniform for both isoforms (Fig 4G and P).

**Figure 4.**
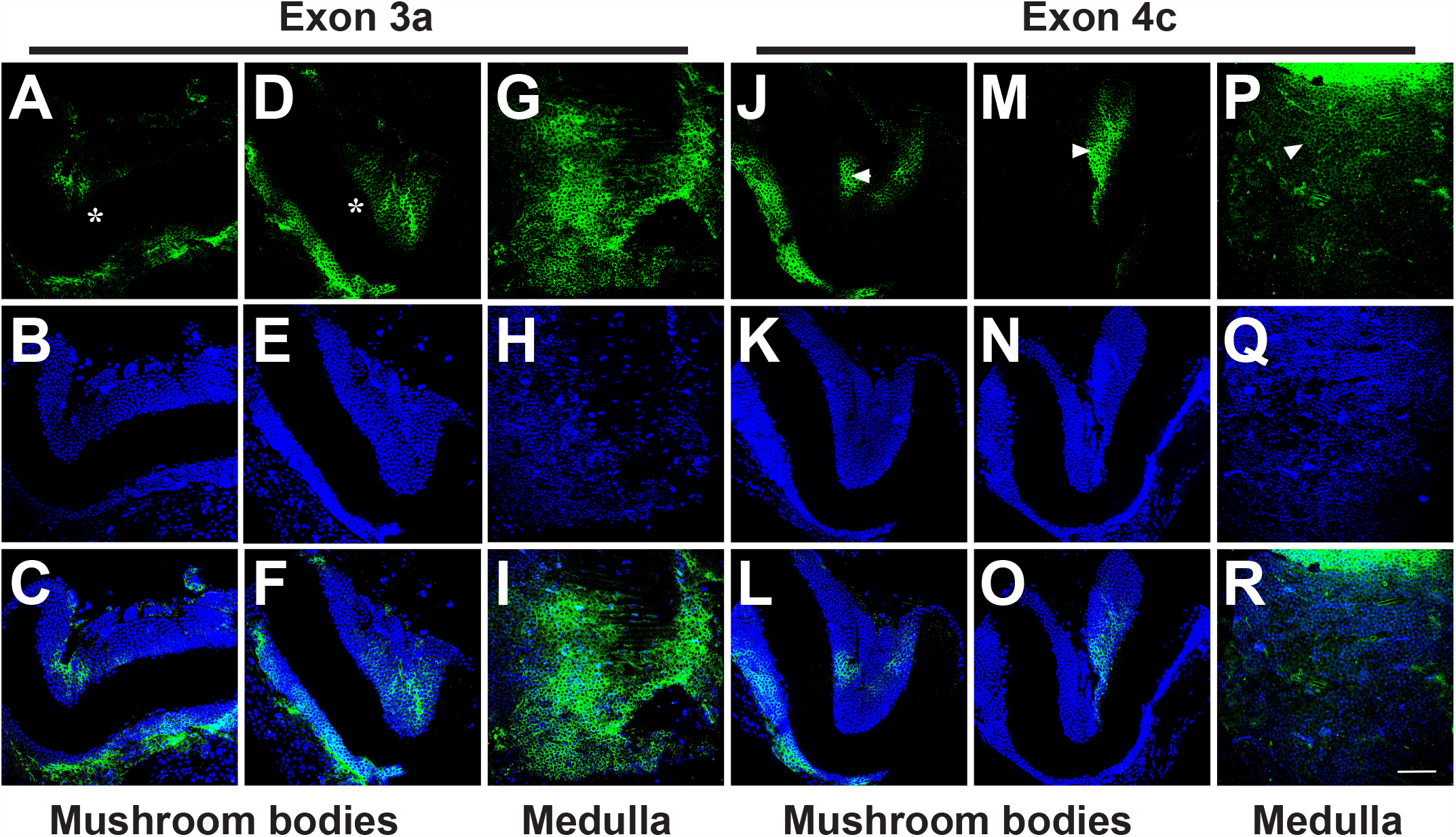
Alternative splicing of *elav* exons 3a and 4c is dynamic in mushroom bodies. Representative RNA in situ hybridizations in worker bees against *elav* exon 3a (A, D and G) and exon 4c (J, M and P) in mushroom bodies (A, D, J and M) and the medulla (G and P) counterstained with DAPI to visualize nuclei (B, E, H, K, N and Q) and merged pictures (C, F, I, L, O and R). Scale bar in R is 30 µm.

### ELAV protein levels are dynamic in mushroom bodies of worker bees

ELAV family proteins are pan-neurally expressed in *Drosophila*. Their expression seems not to be dynamic as judged form antibody stainings, but changes in nuclear and cytoplasmic distributions have been observed ^24^. Consistent with its nuclear localization in flies, ELAV is also mostly nuclear and broadly expressed in about 75% of worker bee brains analyzed (Fig 5A). In the remaining 25%, however, ELAV expression was very dynamic, showing patches of nuclear and cytoplasmic localization, but also small patches of cells with no ELAV expression (Fig 5D,G, J). Because these analyses were done on animals whose previous experience in the field could not be controlled, we wondered whether such localized changes in ELAV expression might be indicative of experience-dependent plasticity.

**Figure 5.**
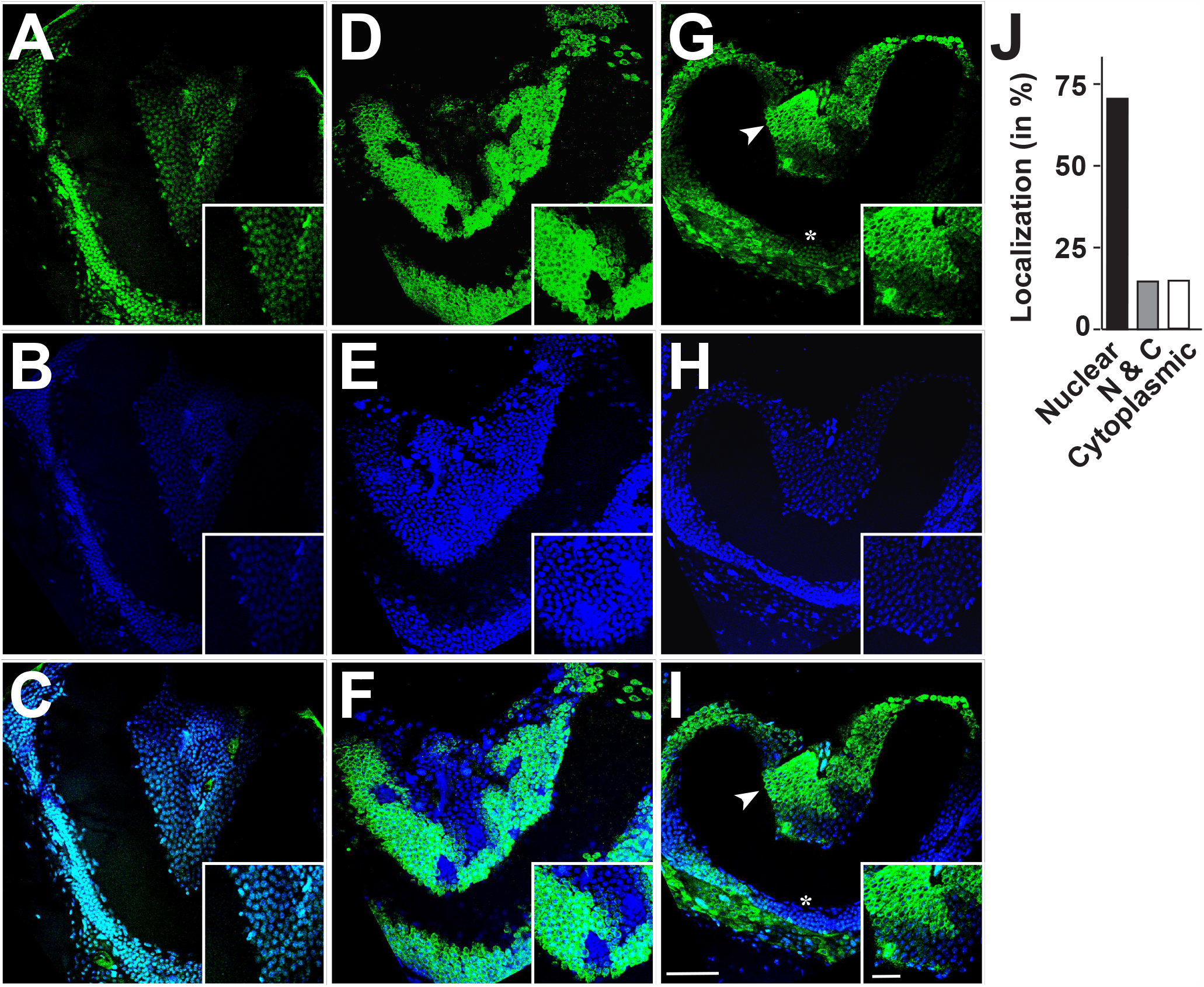
Localization and expression levels of ELAV are dynamic in mushroom bodies. Representative anti-ELAV antibody stainings in worker bees of mushbodies (A, D and G) counterstained with DAPI to visualize nuclei (B, E and H) and merged pictures (C, F and I). Insets show higher magnifications of ubiquitous expression and nuclear localization (inset in A-C), patchy expression with mostly cytoplasmic localization (inset in D-F) and patchy expression with nuclear and cytoplasmic expression (inset in G-I, arrowhead in G and I). The asterisk in G and I indicates nuclear localization in the lower part of the mushroom body. A summary of ELAV localization in mushroom bodies of worker bees is shown in J, n=20. Scale bars are 30 µm in I and 6 µm in the inset.

### ELAV expression and alternative splicing is altered upon learning

Since bees depend on learning and memory to forage, the pronounced loss of ELAV expression in some of the brains of worker bees might reflect inter-individual learning/memory variations. Thus, we thought of testing if such local down regulation might be indicative of a particular individual learning/memory status. To increase the sensitivity of our follow-up molecular analysis we took advantage of the diversity in speed of learning observed among individuals during a 5-trial training by splitting trained bees into fast and slow learners, e.g. bees that responded in the first two trials and every time after the initial response vs bees with a lack of response in the first two trials or with gaps after the initial response (Fig 6A). We then monitored *elav* expression levels from their brains by qPCR at various time points after training (Fig 6B). This analysis indeed revealed that *elav* steady-state mRNA levels had significantly dropped two hours after training in the fast learners compared to slow learners (p=0.13). We therefore thought to analyze alternative splicing of *elav* exons 3a and 4c two hours after training. Indeed, we detected a significant increase in inclusion of exons 3a and 4c in the mushroom bodies, but not in the medulla one hour after training in fast learners (Fig 6C-E). We also analyzed the alternative splicing pattern of ELAV on denaturing polyacrylamide gels, but no differences were detected after learning in this assay, likely because the observed changes occurred only in relatively few cells (Supplemental Fig 4).

**Figure 6.**
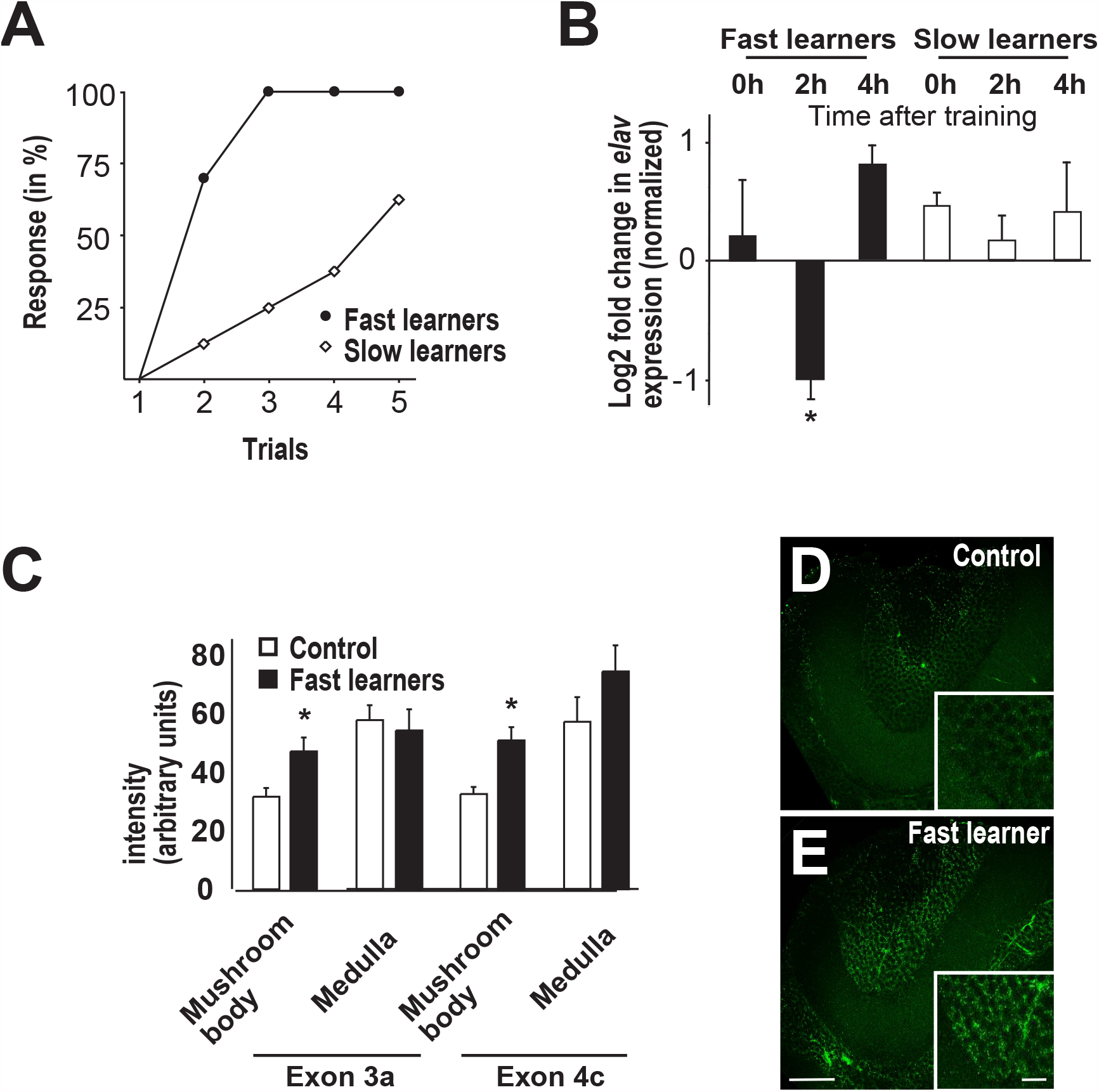
*elav* alternative splicing and expression levels change upon learning. (A) Learning response of bees grouped into fast and slow learners (fast learners n=10 and slow learners n=8). (B) *elav* mRNA expression levels determined by qPCR 0h, 2h and 4h after training shown as mean with the standard error from two to four replicate central brains normalized to *Appl* expression (p=0.13). (C) Inclusion levels of *elav* exons 3a and 4c quantified from RNA in situ hybridizations from control (unpaired) and fast learning (paired) bees in the mushroom body and the medulla (p<0.05, n=7). (D and E) Representative RNA in situ hybridizations against exon 3c in mushroom bodies of control (D) and fast learning (E) bees 2h after training. Scale bars are 30 µm in E and 6 µm in the inset.

## Discussion

Many RNA binding proteins including neuronal ELAV/Hu RBPs are comprised of families of highly related proteins ^16,48^. In case of ELAV family RBPs, they have unique individual functions, but depending on cellular localization and concentrations they can cross-regulate targets making the study of their individual functions difficult ^24,35,36,43^. Therefore we took advantage of honeybees due to presence of only a single *elav* gene to examine whether ELAV is required for learning and memory.

### A role for ELAV in learning and memory

Although neuronal ELAV/Hu family proteins are broadly expressed in the brain, mutants of individual genes in mice and *Drosophila* revealed only subtle developmental defects thus pointing towards a primary role in regulating neuronal functions as e.g. operating in learning and memory ^18,24,49,50^. A knock-out of HuC in mice revealed a role in the synthesis of the neurotransmitter glutamate resulting in reduced neuronal excitability and impaired motor function ^18^. For HuD, roles in learning and memory have been suggested due to its involvement in regulating GAP43 expression which has established roles in learning and memory ^51-53^. Here, overexpression of HuD, which is cytoplasmic, leads to increased GAP-43 expression by increasing mRNA stability. Since in bees *elav* steady state mRNA levels drop for a short period after training early during memory consolidation, this might reflect functional compartmentalization of ELAV/Hu family proteins between nucleus and cytoplasm as bee ELAV is mostly nuclear compared to HuD, which is mostly cytoplasmic in a learning context in the mouse hippocampus ^52^.

The changes in *elav* expression in the brain occur within two hours following learning consistent with a role in memory consolidation. Indeed, our learning protocol was designed to trigger the formation of stable long-term memories, which can be detected several days later ^54^. Such memories are formed through a consolidation process initiated before the end of training and within a few hours, which depends on gene transcription ^10,39^. It is therefore conceivable, that altered levels of ELAV will impact on newly transcribed genes. In particular, expression of ELAV has been linked to implementing splicing programs governing neuronal characteristics such as changes in cell adhesion. Potentially, reduction of *elav* levels could reduce cell adhesion for facilitating creation or pruning of new synaptic connections. Indeed, changes in connectivity, particularly in the mushroom bodies, is an important process underlying long-term memory formation ^55^. Such role is well in agreement with our observations in *Drosophila*, where reducing alternative splicing of the ELAV target *ewg*, a transcription factor, results in increased growth of synapses at the NMJ ^17,56^. Likewise, we observed changes in ELAV alternative splicing in bees leading to an increase in exon 3a and 4c inclusion, which is anticipated to have profound effects on target mRNA binding. In addition, skipping of exon 4, 4a or 4b leads to a frameshift and an altered structure of the third RRM, which will alter target specificity and/or reduce binding affinity.

### Local and dynamic expression changes of ELAV as a hallmark for its role in learning and memory

A hallmark of memory formation is altered local gene expression followed by local changes of neuronal properties and establishment of new connections ^1,3^. Activity induced expression of IEG transcription factors has been associated with memory ^4,5^. Intriguingly, expression of HuD can be induced by neuronal activity ^57^. Stabilization of C/EBP by apELAV1 in *Aplysia* accompanies long-term memory ^58^, although apELAV1 is mainly nuclear in contrast to apELAV2, which is also cytoplasmic ^59^.

In agreement with a role for ELAV in memory formation we find variable expression patterns for ELAV in the mushroom bodies of worker bees typical of IEGs. Even more compelling, the expression pattern of ELAV in the mushroom bodies of worker bees is unique and differs between individuals. Similarly, inclusion levels of alternative exons 3a and 4c also show unique patterns in each individual bee. This can be understood as possible consequences of differences in the previous experience that individuals had had, either within the hive or outdoors (e.g. social interactions, environmental stimuli). Indeed, the mushroom body connectivity is shaped by individual experience during a continuous maturation process ^60,61^. Yet, molecular tools are currently lacking in the honey bee in order to identify those neurons where *elav* expression varies and compare establish interindividual comparisons. In addition, ELAV’s cellular localization also varied in individual cells in the mushroom bodies from nuclear to cytoplasmic. Such differences in cellular localization are expected since *Drosophila* ELAV localizes mostly to the nucleus, RBP9 is cytoplasmic and FNE is found in both compartments. However, ELAV/Hu family proteins also shuttle between nucleus and cytoplasm ^24,62^. Upon removal of ELAV in *Drosophila*, alternatively spliced microexon 4c is included in FNE leading to nuclear localization and regulation of alternative splicing of genes that are otherwise ELAV targets ^35,36^ suggesting a complex network of interactions among ELAV/Hu proteins.

Alternative splicing could serve as an adaptive mechanism to changes in perception, but also to environmental conditions such as toxic insult ^63^. Although learning and memory is affected by neonicotinoids in insects, we did not find any changes in *elav* alternative splicing ^44,64,65^. Since we also could not detect any alternative splicing changes after learning in mRNA from central brains, drastic changes in alternative splicing relevant to learning and memory might occur only in few cells.

### Alternative splicing of a microexon in ELAV/Hu proteins is evolutionary ancient

Human HuB-D genes contain an alternatively spliced microexon in the hinge region between the second and third RRM ^20,66,67^. Intriguingly, we identified an alternatively spliced exon at the same position in the single bee *elav* gene ^44^. Comparison of the sequence between human and insects of this exon shows a high sequence similarity indicating that this exon is evolutionary ancient. Previously, exon duplication between humans and *Drosophila* has been documented in few ion channel genes leading to alternatively spliced exons, but with a different sequence and no longer exons are evolutionary conserved between invertebrates and vertebrates ^46,47^. Intriguingly, ELAV in *Drosophila* has lost its introns due to retrotransposition, but retained microexon 4c ^19^. This microexon is involved in regulating nuclear localization of ELAV and FNE in Drosophila ^35,36,45^, and also affects localization of HuD in human cells ^66,67^. Its increased expression shortly after training thus coincides with an initial nuclear role of ELAV at the memory consolidation phase, which requires transcription ^10,39^.

For most neuronally alternative spliced microexons in mice, Srrm4 is required for their inclusion ^68^. Srrm4 contains a novel evolutionary conserved protein domain ‘enhancer of microexons’ (eMIC) that is present in *Drosophila* Srrm2/3/4 and required for exon inclusion in the *Dscam* exon 9 cluster ^69^. Taken together, a conserved neuronal microexon program is present in vertebrates and insects ^47^.

### Alternative splicing in bee ELAV is confined to unstructured linker regions, but not RNA recognition domains

A main question arising from the presence of multiple highly related genes is whether they act in an overlapping manner. In case of *Drosophila* ELAV family members ELAV, FNE and RBP9 in *Drosophil*a, distinct mutant phenotypes and the lack of major genetic interactions among them suggests largely independent functions ^24^. However, cross-regulation between FNE and RBP9 is present in the regulation of synapse numbers. Likewise, expression in non-neuronal cells or swapping of expression and localization regulatory regions can to a large degree substitute for their individual functions and they can cross-regulate. Overlapping functions even extend to more distantly related Sex lethal (Sxl), which is required for neuronal functions in *Diptera*, but has been recruited in *Drosophila* for sex determination and dosage compensation ^70^. Here, RBP9 is required for maternal inhibition of dosage compensation, a function that is taken over entirely by Sxl during embryogenesis ^24^.

These facts point out that the main distinction among ELAV family members only minimally occurs at the level of RNA recognition. Hence, it is conceivable, that the ELAV family in bees has “merged back” into a single copy gene by incorporating the variable parts between family members by alternative splicing. In this respect, it is very interesting that alternative splicing in bee ELAV occurs in unstructured linker regions between RRMs. It is conceivable, that these regions mediate protein-protein interactions leading to sub-functionalization. Accordingly, the conserved microexon present in the hinge region likely serves such purpose, but the interacting proteins remain to be identified.

Mis-regulation of microexons has been found as a major cause of autism spectrum disorders revealing essential functions for such microexons in neurons ^68^. Notably, inclusion levels of this microexon in bees is altered upon learning and memory formation. Hence, lack of dynamic inclusion of microexons in ELAV/Hu family proteins might point towards a role in establishing the extensive memories often associated with some autism spectrum disorders ^71^.

## Materials and Methods

### Honey bees and treatment

Honey bees (*Apis mellifera*) were collected from flowers or local bee hives in the UK for molecular biology experiments (worker bees were used unless otherwise specified). For behavioral experiments, workers were taken from the experimental apiary on the university campus in Toulouse (France), on the morning of each experiment. Following cold-anesthesia, they were harnessed in metal tubes leaving access to the head, fed with 5 µl of sucrose solution (50% weight/weight in water) and then kept in the dark at room temperature until needed. They were fed in the same way on every morning and evening during the time of each experiment.

### Behavioral assays

Learning and memory capacities were assessed using a standard protocol based on the olfactory conditioning of the proboscis extension response (PER) ^72^, which consisted of 3 learning trials (unless specified otherwise) where animals were trained individually to associate an odorant with a sucrose reward as detailed below. Memory of the association was tested either one hour (short-term memory) or 48 hour (long-term memory) after the last learning trial. In all experiments, bees of both treatment groups were trained in parallel. Each learning trial (40 s) started when the restrained bee was placed in front of an odorless air flow. After 15 s, the setup allowed to deliver an odor (conditioned stimulus, CS) for 4 s by partially diverting the flow in a syringe containing a filter paper soaked with 4 µl of pure odorant. (1-hexanol and 1-nonanol were used, alternatively for different bees; data were pooled after checking for any significant effect of the odorant used). Sucrose (unconditioned stimulus, US: same solution as for feeding) was delivered to the antennae using a toothpick, 3 s after CS onset, for 3 s. This triggered the bee’s reflex extension of the proboscis to lick the reward. Whenever the animal already responded to the CS (conditioned response), it was directly allowed to feed upon US onset. Successive learning trials were separated by 10-min intervals to facilitate memory consolidation ^54^. Memory was assessed by placing the animals again in the conditioning setup, and by presenting them the CS without the US ^72^. The presence or absence of a conditioned response was recorded. In case of no response, sucrose was applied to the antennae at the end of the test, to control for the intact motor response. Bees failing to show an intact reflex were discarded. Bees that responded to the training in the first two trials and that responded every time were classified as fast learners. Bees that responded only two times in the four trials were classified as slow learners. The sucrose and odorants were purchased from Sigma-Aldrich (France).

### Recombinant DNA technology, RT-PCR, qPCR and analysis of alternative splicing

Recombinant DNA technology was done according to standard procedures as described ^73^. Bee *elav* was amplified from oligo dT primed cDNA made from larval brains using primers elav F1 (GCCGCCGGCGCGAACGGAATGGACACAGTCGTACAAC) and elav R1 (GCGTCTAGAGGCGCGCCTCTACGCCGCCTTGCTCTTGTTCGTCTTGAAGC) and cloned into a modified pBS SK+ using NgoMIV and XbaI. Clones (n=45) were sequenced using primers elav F1 and elav R1. RNA extraction from whole bees or dissected bee brains and RT-PCR was done as described ^74^. *elav* expression at different timepoints was compared to *Appl* expression using primers AM elav qF3 (CCCTCTTCTCGAGCATTGGCGAGGTTG) and AM elav qR3 (GCCGTACGGGCTGAATAGATTCTCCAG) to amplify the constant part of *elav* and normalized to unpaired control animals using qPCR as described ^65^. For high resolution analysis of *elav* alternative splicing primers elav F2 (GTCGCGGATACTTTGCGACAACATCAC) and elav R2 (CCCGGGTAGCATCGAGTTTGCCAATAGATC) were used to amplify *elav* from cDNA. One of the primers was labeled using gamma^32^P-ATP (NEN) and PCR products were separated on sequencing type denaturing polyacrylamide gels. Polyacrylamide gels were dried, exposed to phosphoimager screens (BioRad) and quantified with QuantityOne (BioRad).

### RNAi, Western analysis, RNA in situ, antibody stainings and imaging

For RNAi knockdown in bees, *elav* and GFP DNA templates for *in vitro* transcription were amplified for *elav* from a cloned cDNA with primers *Apis* ELAV T7 RNAi F1 (GGAGCTAATACGACTCACTATAGGGAGAATGATGGCGAACGGAATGGACACAG) and *Apis* ELAV T7 RNAi R1 (GGAGCTAATACGACTCACTATAGGGAGACTACGCCGCCTTGCTCTTGTTCGTCTTG) and for *GFP* a 700 bp fragment was amplified with primers GFP T7 RNAi F1 (GGAGCTAATACGACTCACTATAGGGAGACTGTTCACCGGGGTGGTGCCCATC) and GFP T7 RNAi R1 (GGAGCTAATACGACTCACTATAGGGAGACTTGTACAGCTCGTCCATGCCGAGAG).

Double stranded RNA was generated by *in vitro* transcription with T7 polymerase with the MegaScript kit (Ambion) for 3 h according to the manufacturer’s instructions. After digestion of the template with TurboDNAse (Ambion), dsRNA was phenol/chloroform extracted, ethanol precipitated and taken up in RNAse free water at a concentration of 5 µg/µl. The dsRNA (250 nl) was then injected into the brain through the median ocellus with a Nanoject II microinjector (Drummond). RNAi efficiency testing for ELAV was done from dissected central brains by Western blotting according to standard protocols as described ^73^ using a polyclonal rat anti-ELAV antibody generated against Drosophila ELAV (1:800) ^45^ and secondary HRP-coupled goat anti-rat antibody (1:5000, GE Healthcare) by chemoluminescence detection (Pierce) according to the manufacturer’s instructions.

Brain antibody stainings were done with rat polyclonal anti-ELAV antibody (1:200) as described ^17^ and counterstained with DAPI (1 µg/ml).

To make probes for RNA in situ hybridizations, a pBS SK+ vector was modified by cloning a U-rich stem loop at the end of the in vitro transcript using EcoRI and KpnI and phosphorylated and annealed oligos RNA IS stem 1A (CTCAAACACATATATACATATACATATAGGGGTACATACATATATACATATATACTC GAG) and B (AGTATATATGTATATATGTATGTACCCCTATATGTATATGTATATATGTGTTTGAGG TAC), tango13A (TATGGTAAGCCAGATGCATGGGTGCAGGACAACACGTCGAAGttaGATATCG) and B (AATTCGATATCtaaCTTCGACGTGTTGTCCTGCACCCATGCATCTGGCTTACCATACTC G). ELAV alternative exons were then cloned with XhoI and PstI using phosphorylated and annealed oligos apis elav 3a A (TCGAGACAACATCACCGTACGACAGTTTGTGACCGGCGGCGGAGACTATTTGCCCGGATTGTCGAAAAGTACTGAATTCCTGCA) and B (GGAATTCAGTACTTTTCGACAATCCGGGCAAATAGTCTCCGCCGCCGGTCACAAACTGTCGTACGGTGATGTTGTC) and apis elav 4c A (TCGAGGCCGCTTCAGCACTGGCAAGGCCATGCTTGCCATTAACAAAGGCTTACAGAGGTACAGCCCGCAGTACTGAATTCCTGCA) and B (GGAATTCAGTACTGCGGGCTGTACCTCTGTAAGCCTTTGTTAATGGCAAGCATGGCCTTGCCAGTGCTGAAGCGGCC). Vectors were linearized with Acc56I and DIG-dUTP (Roche) labeled anti-sense transcripts were generated by in vitro transcription with T3 RNA polymerase (Ambion) in 10 µl from 1 µg template DNA. These transcripts were cleaned by centrifugation through a G50 Microspin column (GE Healthcare) in a final volume of 50 µl. In situ hybridizations were done on whole brains in 50% formamide buffer as described ^17^ using 1:500 diluted probes at 39 °C for 3d and washed overnight in hybridization buffer. DIG-labeled probes were then visualized with an FITC conjugated anti-DIG antibody (Roche) and counterstained with DAPI.

For brain imaging, confocal Z stacks were taken using a Leica SP5/SP2, using a 40x-oil objective. For the quantification of stained Kenyon cells, the cross-section equal to the width of the calyx was scanned and the fluorescence intensity quantification was performed as previously described using ImageJ ^24^. For the imaging of the calyces of the mushroom bodies of honey bee brains, single optical sections were taken in the x-y plane. The image acquisition settings were kept identical for all preparations.

### Protein modeling and statistics

Structure modeling was done using Phyre2 of individual RRMs and assembled into a composite of all three RRMs ^75^. Unstructured loop parts in RRMs and the hinge region were inserted manually.

Multiple planned pairwise comparisons of expression levels were done by ANOVA followed by Fisher’s protected least significance difference post-hoc test using StatView. To compare proportions of conditioned responses between groups, a repeated-measure analysis of variance (ANOVA) was run for the acquisition data (one factor, *treatment*, with *trial* as the repeated measure), and a simple ANOVA for retention data ^76^. Post-hoc comparisons of rates of CS-specific responses were done using the Fisher’s exact test.

## Acknowledgments

We thank the Winterbourne garden (Birmingham) and Lucie Hotier (Toulouse) for providing bees, Roland Arnold and Reinhard Stöger for discussions and comments on the manuscript. For this work we acknowledge funding from the Sukran Sinan Fund, the Genetics Society, the Biochemical Society and BBSRC. JMD acknowledges funding from the CNRS and Université Paul Sabatier.

## Authors’ contribution

PU, IUH and MS performed molecular biology experiments, PU, JKG and ND performed behavioral experiments, and JKG performed antibody stainings and in situ hybridization experiments. JMD designed behavioral experiments. MS conceived the project and wrote the original draft of the manuscript. JMD, IUH and all other authors reviewed and edited. M.S. and JMD supervised and acquired funding.

## Data availability

All data are available in the main text or the supplementary material.

## Figure legends

**Supplemental Figure 1**. Sequence comparison of ELAV family proteins of bees and *Drosophila*.

(A) Alignment of single *Apis mellifera* ELAV with the three *Drosophila melanogaster* orthologues ELAV, FNE and RBP9. Long and short lines on top indicate the three RRMs and RNP motives, respectively. Black triangles depict intron positions. Alternative exons 3a, 4a, 4c and 4d are indicated at relevant intron positions. The hatched line indicates the sequence deleted in exon 4b by alternative splicing. Note that skipping of exons 4, 4a or 4b will result in a truncated protein due to a frameshift (fs).

(B) Phylogenetic tree of *Apis* and *Drosophila* ELAV family proteins.

**Supplemental Figure 2**. Structural models of bee ELAV depicting the position of alternative exons.

(A) Full length ELAV. Note that alternative exons are in flexible linker regions in RRM2 (exons 3a and 4a) and the hinge region (exons 4b, 4c and 4d).

(B) Truncated ELAV resulting from skipping of exon 4.

(C) Truncated ELAV resulting from inclusion of exon 3a and skipping of exon 4. Note that exon 3a adds an additional B-sheet to the truncated second RRM potentially increasing RNA binding.

(D) Truncated ELAV resulting from inclusion of exons 3a and 4C, and skipping of exon 4. Note that exon 4c adds a second alpha-helix to the truncated second RRM.

**Supplemental Figure 3**. Representative gels from the developmental analysis of ELAV alternative splicing quantified in Fig 3G-I.

(A and B) Top and bottom gel part of a representative 5 % denaturing polyacrylamide gel separating ^32^P-labeled alternative splice products from indicated developmental stages on top.

Length of PCR products from splice variants are indicated at the right. # The 120 nt product can be either 3, 4c, 5 or 3, 3a, 4d, 5. M: Marker.

(C and D) Analysis of alternative splicing from indicated developmental stages proximal (*Kpn*I) and distal (*Fok*I) of exon 4 from ^32^P-labeled labeled forward (C) or return (D) primer after digestion with either *Kpn*I or *Fok*I on a representative 5 % denaturing polyacrylamide gels. ^ The 123 nt products are either 3, 4c, 4d, 5 or 4b, 4c, 4d, 5. ^**^**^ The 120 nt products are either 3, 3a, 4d, 5 or 4b, 4c, 5 or 3, 4c, 5. * The 78/80 nt products are either 4b, 5 or 3, 5 +/- 4d. M: Marker.

**Supplemental Figure 4**. Representative gels of ELAV alternative splicing after learning. (A and B) Top and bottom gel part of a representative 5 % denaturing polyacrylamide gel separating ^32^P-labeled alternative splice products from naïve and trained bees from unpaired and paired conditioning of conditioned (sugar) and unconditioned (odor) stimuli at the indicated time after training. Length of PCR products from splice variants are indicated at the right. # The 120 nt product can be either 3, 4c, 5 or 3, 3a, 4d, 5. M: Marker.

(C and D) Analysis of alternative splicing from naïve and trained bees from unpaired and paired conditioning of conditioned (sugar) and unconditioned (odor) stimuli at the indicated time after training proximal (*Kpn*I) and distal (*Fok*I) of exon 4 from ^32^P-labeled labeled forward (C) or return (D) primer after digestion with either *Kpn*I or *Fok*I on a representative 5 % denaturing polyacrylamide gels. ^ The 123 nt products are either 3, 4c, 4d, 5 or 4b, 4c, 4d, 5. ^**^**^ The 120 nt products are either 3, 3a, 4d, 5 or 4b, 4c, 5 or 3, 4c, 5. * The 78/80 nt products are either 4b, 5 or 3, 5 +/- 4d. M: Marker.

