## Supplementary figures and images for "Dynamically expressed ELAV is required for learning and memory in bees"

### Supplemental Figs 1-4

#### Supplemental Figure 1

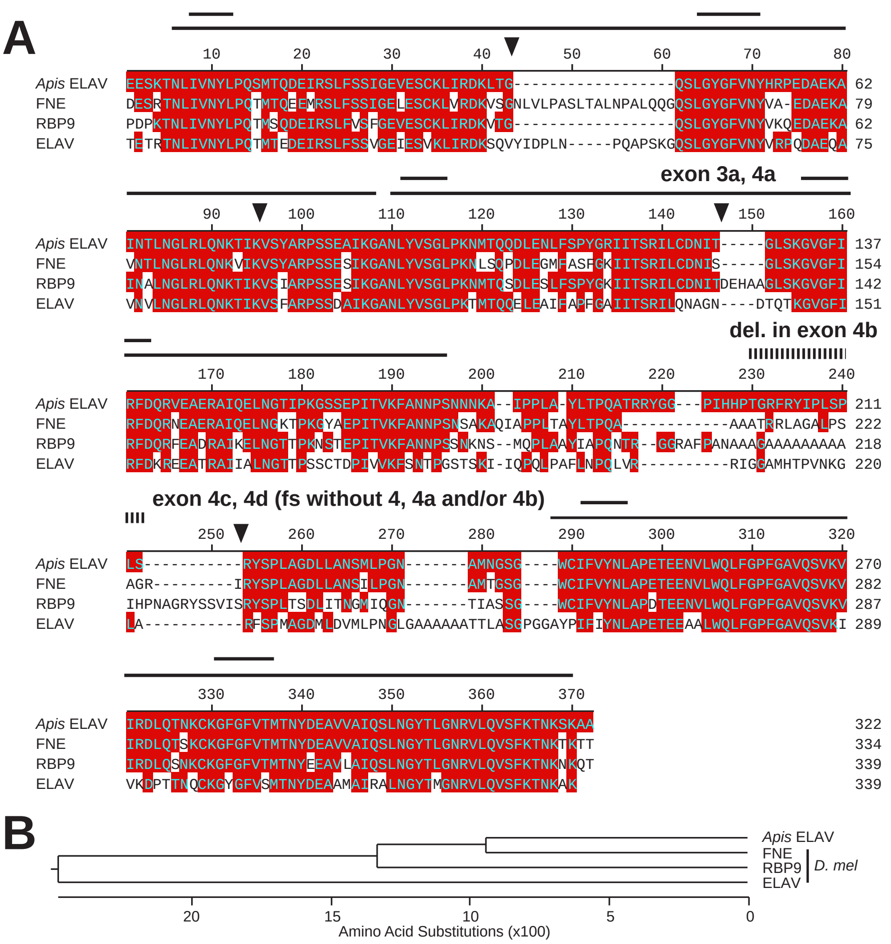

##### Supplemental Figure 2

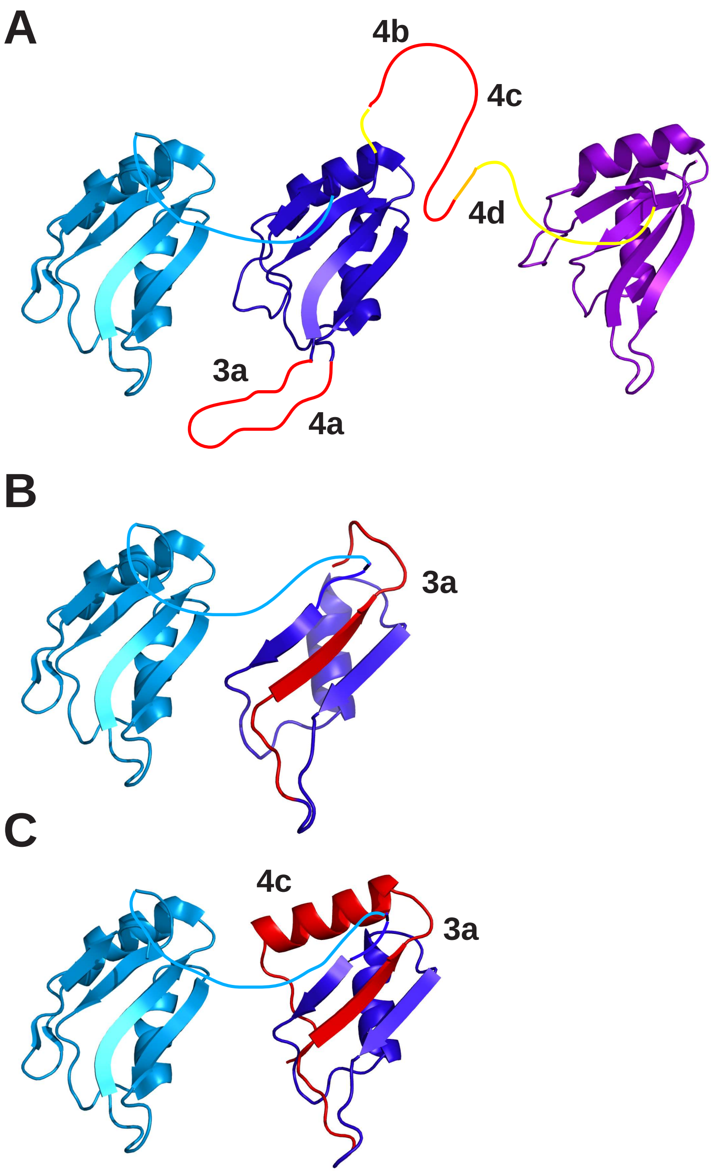

#### Supplemental Figure 3

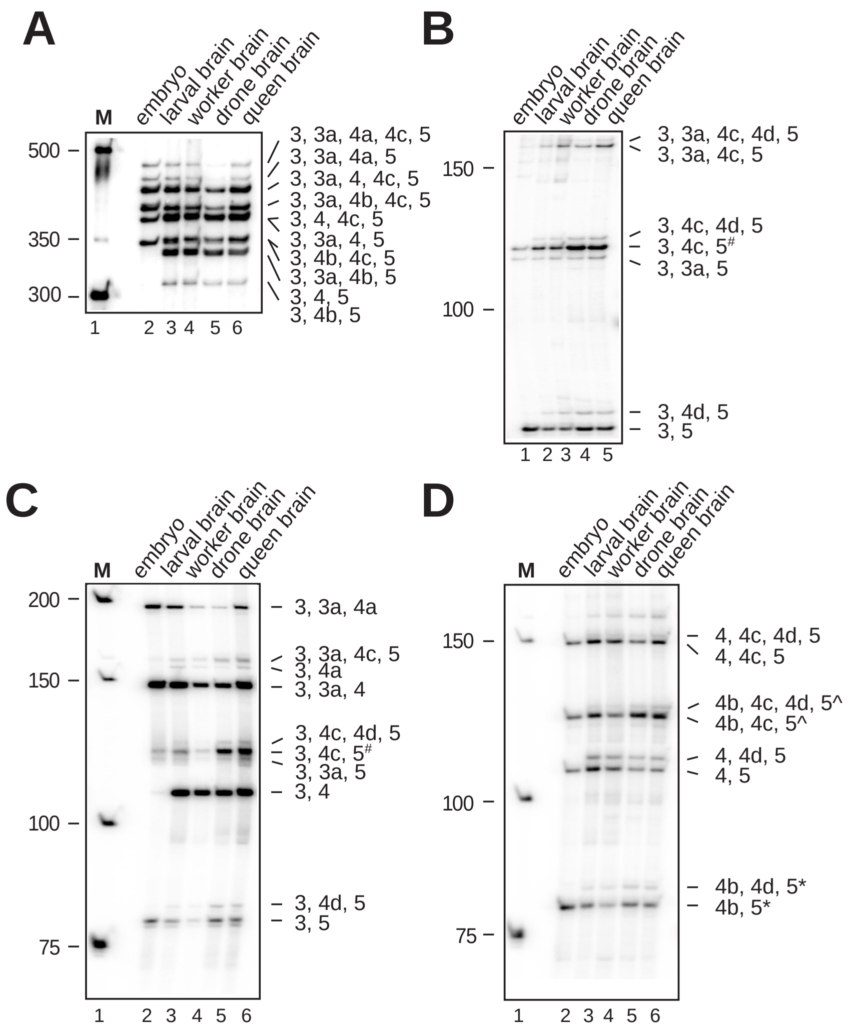

#### Supplemental Figure 4

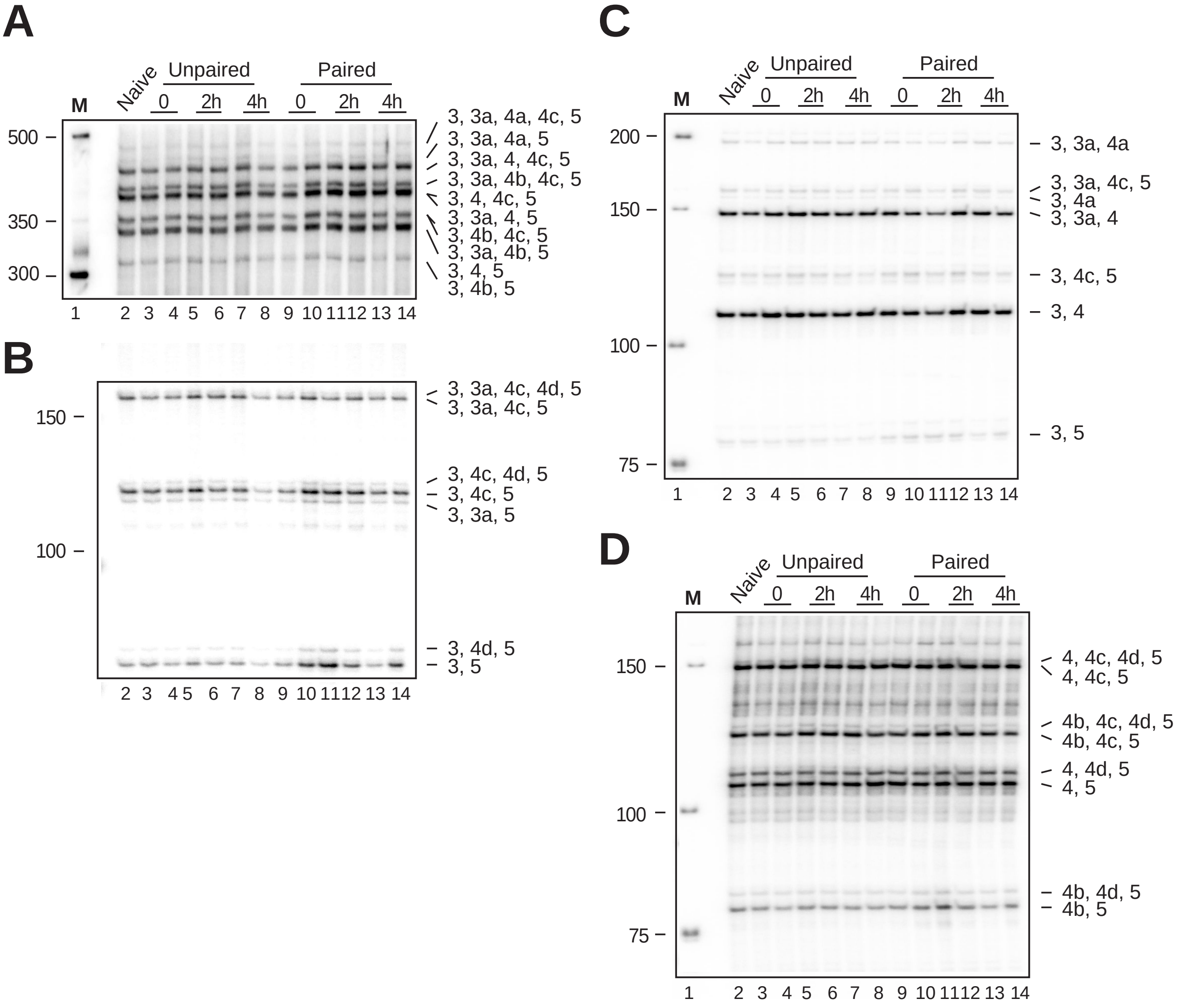
